# Reducing Honey Bee Winter Mortality with Molybdenum Supplementation: Field Evidence Across Europe

**DOI:** 10.1101/2025.07.23.666091

**Authors:** María Benito-Murcia, Clémence Riva, Eduardo José García-Vicente, Ana Pérez, María Martín Domínguez, Noelia Hermosilla, Précillia Cochard, Leonidas Charistos, Fani Hatjina, Mathieu Holmiere, Salomé Martínez-Morcillo, Benjamin Poirot, David Risco, Sébastien Floquet

## Abstract

Nutrition is key to improve honey bees’ resilience to environmental stress factors that threaten their health. Yet, little is known about the micronutritional needs of colonies, in particular about molybdenum, an essential trace element in biology. This study focuses on a coordination complex based on molybdenum Mo(+V) which was assessed for compatibility with standard beekeeping practices. It was found to be non-toxic for bees, stable and easy to use. Importantly, it does not leave any residues in honey produced. This study investigated how supplementation with few milligrams of this complex can improve hive performance at early spring and reduce winter mortality. Over a two-year field campaign, 283 beehives spread over 6 apiaries located in Spain, Greece and France were involved: 142 beehives supplemented with the molybdenum-based compound and 141 beehives as a control group. Supplementation resulted in a significant reduction in winter colony losses, averaging a 44% decrease, ranging between 27% and 75% in the different apiaries. The risk of death was twice lower in the Na-Mo_2_O_4_-EDTA group compared to the control group (risk ratio: 0.51). The impact on bee populations and honey production was also evaluated. A significant increase in honey reserves within the brood area of 107% was observed in the Greek apiary, whereas no comparable effect was detected in Spain, suggesting that local environmental conditions or management practices may influence this parameter. This study highlights the importance of molybdenum in the management of honey bees as an efficient tool to reduce the winter mortality of the colonies.

**Highlights:**

- Molybdenum-based feed supplementation reduced honey bee winter mortality by 44% in average and up to 75%
- There are no adverse effects on honey bee health or honey and wax composition.
- Supplement showed variable effects on honey reserves and no effects on bee population.
- ALP levels increased transiently, while the TAC was unaffected.Supplementation with Molybdenum-based compound represents a promising strategy to enhance colony overwintering success

## 1. Introduction

The managed honey bee (*Apis mellifera*) play a key role in pollination and are essential for both plant biodiversity and food supply. The vast majority of flowering plants—including around 75% of major commercial crops and 35% of the global food supply—depend at least partly on pollinators, with honey bees playing a crucial role (Aizen et al., 2009; Klein et al., 2007; Ollerton et al., 2011). Beyond pollination services, beekeeping is an important economic sector that produces honey and other valuable beehive products, such as propolis and beeswax, and it is of great interest to enhance rural livelihoods (Yap and Devlin, 2015).

Over the last 30 years, severe colony losses, particularly during winter, have been reported across continents (Bruckner et al., 2023; Gray et al., 2023; Requier et al., 2018; vanEngelsdorp and Meixner, 2010). Given the economic and ecological value of honey bees, colony losses have raised significant concerns among the scientific community, policymakers, and the public. It is well established that various environmental stressors are contributing to honey bee mortality (Goulson et al., 2015; Hristov et al., 2020; Staveley et al., 2014), including diseases and parasites such as the mite *Varroa destructor* (Hernandez et al., 2022), landscape alteration (Vanbergen, 2014), beekeeping management practices (Jacques et al., 2017), nutritional shortages (Naug, 2009; Tsuruda et al., 2021) and climate changes which can affect not only managed honey bees (Le Conte and Navajas, 2008) but also other unmanaged insects (Abbasi, 2025). Consequently, up to 30% of hives are lost annually in Europe (Gray et al., 2023; Laurent et al., 2016). For these reasons, the development of new solutions to strengthen honey bee health and resilience to stress is of great interest.

Quality nutrition is one way to improve honey bee health, reduce mortality (Alaux et al., 2010; Dolezal et al., 2019) and mitigate the effects of other stressors (Castle et al., 2022), while nutritional limitation can lead to increased disease susceptibility (Dolezal and Toth, 2018). However, optimizing bee nutrition is complex. It requires a perfect balance between macronutrients and micronutrients, which vary throughout the seasons, climate conditions, or available resources (Bonoan et al., 2018, 2017; Lau et al., 2023).

To overcome nutritional limitations, beekeepers rely on feed supplements given during dearth periods, usually in the form of sugar syrup and fondants. These nutritional supplements primarily deliver carbohydrates to the colony, with some formulations also containing proteins (Pavlović et al., 2025; Ricigliano et al., 2022).

Micronutrients include vitamins and minerals. Among minerals, trace elements (TE), serve a wide variety of functions in the biochemical and physiological processes, and are required in small quantities. While the link between macronutrition and the health of honey bee colonies has been studied many times (Brodschneider and Crailsheim, 2010; Tsuruda et al., 2021), studies on the importance of TE for honey bees are scarce (Herbert, 1979; Zhang et al., 2015). Essential TE cannot be produced by bees and the needs of honey bees are met by pollen and, to a smaller extent, by turbid waters (Bonoan et al., 2017). However, the chemical composition of pollen varies among flower species (Bay et al., 2021) and the availability of micronutrients for honey bees differs across space and time.

Among TE, molybdenum (Mo) plays an essential role in insects (Dow, 2017). It is a cofactor of a wide variety of enzymes which have a critical role in the catalysis of redox reactions (Hille, 2002; Schwarz et al., 2009). In eukaryotes, these enzymes mainly belong to the Sulfite Oxidase (SO) and Xanthine Oxidase (XO) families (Peng et al., 2018), which are useful for the degradation of xenobiotics with a broad substrate spectrum. For bees, the main source of Mo is pollen. It was found that honey bees naturally contain about 0.4 ppm of Mo on average (Fuior et al. 2025). While potential benefits of Mo-based compounds on honey bee health have been identified (Fuior et al., 2025) the role of molybdenum for honey bee health remains unknown. .

The complex used in this study (Figure 1A) consists of the central cluster [Mo_2_O_4_]^2+^, in which the two Mo atoms are in the reduced +V oxidation state, surrounded by an organic EDTA molecule to form the anionic complex [Mo_2_O_4_(EDTA)]^2-^. The charge is counter-balanced by two cations, which can be Na⁺ to give the complex of formula Na_2_[Mo_2_O_4_(EDTA)], abbreviated hereafter as Na-Mo_2_O_4_-EDTA.

**Figure 1.**
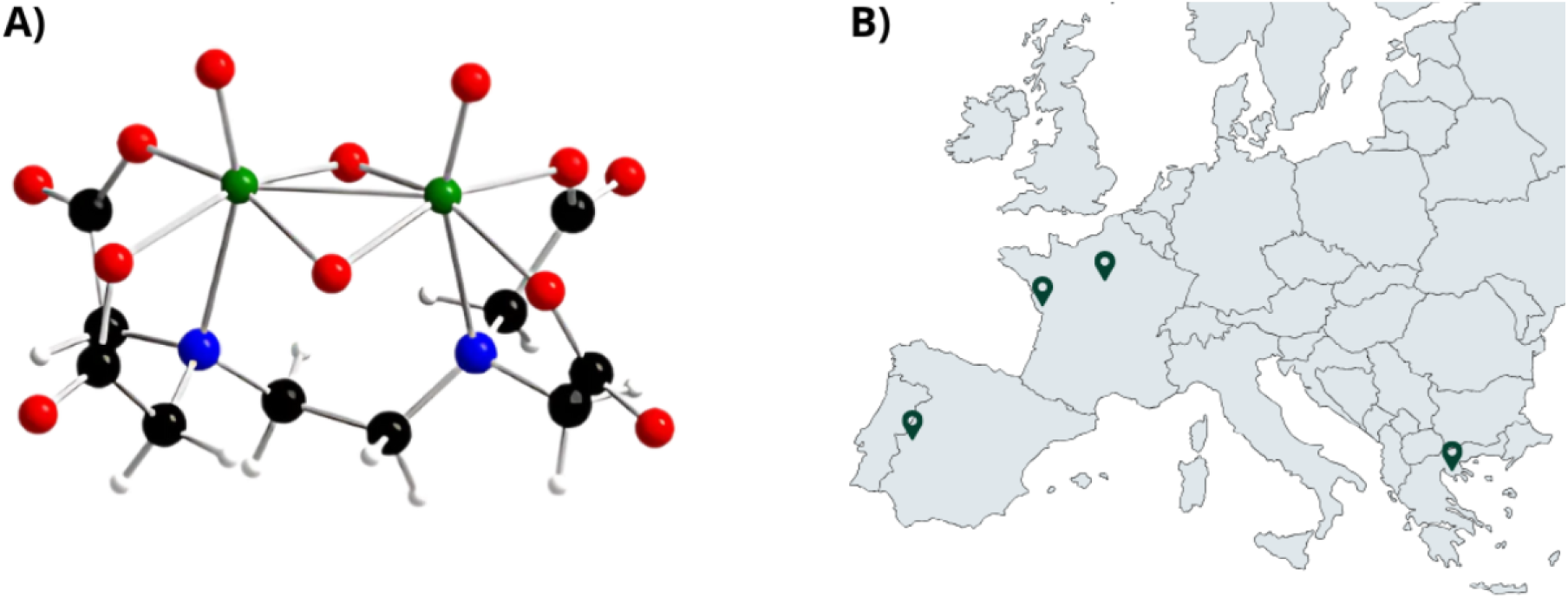
A) chemical structure of the complex [Mo_2_O_4_-EDTA]^2-^, B) map showing the localization of the apiaries (and bee genetics) used in this study in France (A, mellifera, hybrid Buckfast), Greece (A, m, macedonica) and Spain (A. m. iberiensis).

The primary objective of the present study was to evaluate the effects of Na-Mo_2_O_4_-EDTA supplementation on reducing honey bee colony winter mortality and on boosting spring colony dynamics. The second objective was to understand the supplement mode of action, and in particular its antioxidant capacities. Finally, it was aimed to establish whether the compound meets the beekeeping industry standards, in terms of non-toxicity, stability, and its potential impact on honey and wax. All field experiments were done to be as close as possible to European beekeeping common practices. We collected data in six different apiaries in Greece, France, and Spain (see Figure 1B) to test the feed supplement in various climates, with different genetic stocks and beekeeping practices.

## 2. Materials and methods

### 2.1. Synthesis of Na-Mo_2_O_4_-EDTA complex and preparation of syrup

The complex Na-Mo_2_O_4_-EDTA was prepared using a one-pot procedure in water as reported by Fuior et al. (2022). The quality of the complex was checked by routine methods (^1^H NMR, FT-IR, ESI-MS, elemental analysis) before use. By considering an average level of Mo in honey bees of 0.4 ppm (µg/g) (Fuior et al. 2025) and an average weight of 80 mg per bee, a global dose corresponding to 30 ng of Mo per bee was estimated. For a population ranging from 10000 to 60000 honey bees per colony with a lifespan of 40-45 days, we globally estimated the need of the Mo in the 2-4 mg range per colony per season. In the field trials, the quantity of Mo-complex given to the colonies was thus fixed to 8 mg, which corresponds to 2.2 mg of molybdenum. For the syrup preparation, mother aqueous solutions ranging from 100 mg/L to 1.6 g/L of Na-Mo_2_O_4_-EDTA were prepared by dissolving the corresponding mass of complex in 1 L of demineralized water. A small amount of aqueous solution, was then diluted either in commercial beekeeping syrup or a sucrose solution at a concentration of 50% (w/w) to get 2mg of Na-Mo_2_O_4_-EDTA per liter of syrup. The complex is perfectly soluble in these conditions. To reach the dose of 8 mg of complex per hive per feeding, 4L of syrup at 2mg/L is then required for each hive. The concentration in Mo was checked by ICP-MS analyses (Laboratoire de l’Environnement et de l’Alimentation de la Vendée, LEAV in La Roche-sur-Yon, France).

### 2.2 Assessment of the complex Na-Mo_2_O_4_-EDTA stability and its safety for honey production

The stability and safety of the Mo-complex as feed additive was evaluated in accordance with Regulation (EC) No 1831/2003 and EFSA FEEDAP Panel guidance (EFSA Journal 2017;15(5):5021), which among others requires proofs on non-toxicity for target species and beehive products, as well as product stability.

The long-term stability of Na-Mo_2_O_4_-EDTA both in water and sugar syrup was addressed by analysis of Mo content by ICP-MS. ICP-MS analysis of Mo was performed on an aqueous solution of Na-Mo_2_O_4_-EDTA at 1.6 g/L (the solubility of Na-Mo_2_O_4_-EDTA is higher than 1000 g/L in water), and on sugar syrup of the brand “Sirop Royal” containing approximately 2 mg/L of Na-Mo_2_O_4_-EDTA (about 3 µM of complex), following the protocol NF EN 13805/ICP MS. The analysis was performed by the laboratory LEAV in France. The Mo-content was studied for two years with one analysis performed every two months.

One important point for beekeepers and honey consumers is to ensure that honey quality is not affected by the feeding and that the feed supplements are not found in the honey and wax. To address this point, we measured the Mo content of honey and wax and the quality of honey from hives fed with Na-Mo_2_O_4_-EDTA. Eight colonies from an experimental apiary in the West of France were randomly split into two experimental groups. Hives from the supplemented group were first fed in February 2020, with 2.5kg fondant containing sugar and 15% honey, and 24 mg of Na-Mo_2_O_4_-EDTA. Then, in April, they received sucrose syrup (50% w/w) containing 2 mg/L of Na-Mo_2_O_4_-EDTA at Day 0 (D0, 2 liters) and at Day 7 (D7, 2 liters). The control group received the same fondant and syrup at the same time, without Na-Mo_2_O_4_-EDTA supplement. The Mo content of the fondant and syrup was checked by ICP-MS. The total dose of Na-Mo_2_O_4_-EDTA received by each colony was 32 mg (i.e. 8.8 mg of the trace element Mo). Two months later, at the beginning of July 2020, we randomly collected samples of wax and honey in the brood area and in supers in all hives. For each sample, the Mo content was determined by ICP-MS by QSI GmbH laboratory (Bremen, Germany), with a limit of quantification at 0.1 mg/L. EDTA and its main degradation products, that are iminodiacetic acid (IDA), ethylenediaminetriacetic acid (EDTria) and ethylenediaminediacetique acid (EDDA) (see Figure A2, Appendix) were analyzed by GC-MS studies performed by AROMALYSE laboratory (Dijon, France). Additionally, the analysis of honey quality was performed by QSI GmbH laboratory (Bremen, Germany), according to the European Union regulations (pH, humidity, HMF, sugars, conductivity, diastase). For the last one, the honey samples were pooled to have one analysis of honey from the brood area and another one from the super area for control and supplemented groups.

### 2.4 Field experiments

The field trials were conducted in compliance with Directive 2010/63/EU, which exempts invertebrates such as honey bees from mandatory ethical approval. Nevertheless, all procedures strictly followed EFSA guidelines (EFSA Journal 2013;11(7):3295) to ensure animal welfare standards were upheld throughout the study.

To explore the effects of supplementation on the health of colonies in different field scenarios, four experiments including different climatic conditions, beekeeping management practices and seasons of the year, were designed. We carried out these experiments to assess the safety of Na-Mo_2_O_4_-EDTA for bees and the effect of supplementation on winter mortality, strength, and productivity of colonies. For all field experiments, we used queen-right colonies, with no clinical signs of disease, randomly split using the “sample” function in R into a control group that received plain syrup, and a supplemented group receiving syrup with Na-Mo_2_O_4_-EDTA at the concentration of 2mg/L.

#### 2.4.1 Greece 2022 - honey bee mortality, honey surface and bee population

We used 30 colonies of *A. mellifera macedonica* in Langstroth nucleus hives at the Hellenic Agricultural Organization ‘DEMETER’ apiary in Nea Moudania, in Greece. In early May 2022, the 30 colonies were equalized with sister queens of the same age and genetic line, 2 frames of brood, 1 frame of honey, and 1 embossed wax frame. Each hive contained approximately 6000 to 9000 honey bees. The 30 honey bee colonies were randomly assigned to two groups of 15 hives each: a control group and a group receiving supplementation.

The experiment started mid-May 2022, where 15 colonies were fed twice (on D0 and D7) with 2L of supplemented syrup. At the end of both feedings, each colony has received 8 mg of the Na-Mo_2_O_4_-EDTA complex. The remaining 15 colonies were used as control. To confirm the safety of the complex Na-Mo_2_O_4_-EDTA, we assessed honey bee mortality by equipping each hive with a trap at their entrance (Gary, 1960). Dead bees were collected and counted in each trap every 4 days. On Day 56, the cumulated number of dead bees was summarized to one count.

The surface of honey and the adult bee population were evaluated at D0 and D56 by visually assessing the percentage of comb surface covered by honey and by bees, according to the standardized “Liebefeld” method. Each percentage value was then quantified into an actual honey surface or bee population based on the fact that each side of the Langstroth frame has a surface area of 880 cm^2^ and that each 10 cm^2^ contains approximately 12.5 honey bees (Guzman-Novoa et al., 2024).

#### 2.4.2 France 2023 - winter mortality

A total of 93 colonies of *A. mellifera (hybrid Buckfast)* were distributed across two apiaries located in different regions of France: Fontenay-le-Comte (Vendée) and Rochefort-en-Yvelines (Yvelines). In each apiary, hives were randomly assigned to the control or experimental groups. The experiment duration was 90 days, beginning in early November (Day 0) and ending in early February (Day 90). Colonies were fed twice (on Day 0 and Day 7) with 2L of syrup, either plain or supplemented with Na-Mo_2_O_4_-EDTA. All colonies were treated against *V. destructor* in August. Mortality rates were assessed on Day 90, counting dead and alive colonies. Queen right colonies at the time of assessment were considered as surviving ones.

#### 2.4.3 Spain 2023 and 2024 - winter mortality, honey surface and bee population

Two similar experiments were conducted in Spain in 2023 and 2024 using queen right colonies of *A. mellifera iberiensis* housed in Layens-type hives (12 frames). We selected two apiaries located 12 km from each other in Trujillo, Extremadura. The feeding was administered using vertical feeders. Given the impact of *V. destructor* on winter mortality, all colonies were treated against the mite in late summer 2023 and 2024. The surface of honey and the adult bee population were evaluated by visually assessing the percentage of comb surface covered by honey and by bees according to the standardized “Liebefeld” method. The local beekeeper was using Layens-type hives, an horizontal hive model with no super added during the season. So the evaluation of honey surface and number of bees was performed at the all colony level. Each percentage value was then quantified into an actual honey surface or bee population based on the fact that each side of the Layens frame has a surface area of 1500 cm^2^ and that each 10 cm^2^ contains approximately 12.4 honey bees (Guzman-Novoa et al., 2024).

##### a. Spain overwinter 2023

Each apiary consisted of 40 colonies, divided equally into two experimental groups. The study duration was 265 days, from the beginning of November 2023 to the end of July 2024. On Day 0, all colonies were evaluated for honey reserves and bee population. Feeding was conducted on Day 0, Day 15, Day 120, and Day 135. At each of these time points, colonies received 2L of syrup. While both feeding and colony assessments were scheduled on the same day, the feeding was always performed after the parameter assessment. Winter mortality was assessed in early March on Day 120. Additional evaluations of honey reserves and adult bee populations were carried out on Day 120 and Day 265, following the same methodology as previously described. Colonies that died during the experiment were removed from the dataset to avoid a bias with the over-representation of zeros.

We measured the oxidative stress status using total antioxidant capacity (TAC) and the activity of the enzyme alkaline phosphatase (ALP, EC 3.1.3.1). Nurse bees were collected directly from the brood frames to select individuals in close contact with open brood. Immediately after field collection, bees were transported to the laboratory under refrigerated conditions and stored at −80 °C until biomarker analysis. For the determinations of TAC and ALP, homogenates of whole honey bees were prepared. Per triplicate, ten bees were pooled and homogenized (1:10 weight:volume) in ice-cold 40 mM phosphate buffer containing 10 mM NaCl (pH 7.4) and 1% Triton X-100, using an Ultra-Turrax T25 homogenizer at maximum speed. Samples were kept on ice during the entire procedure. The homogenates were centrifuged at 16,000 ×g for 20 minutes at 4 °C, and the resulting supernatants were used for all biochemical assays, using a microplate reader (PowerWave^TM^ HT 340 spectrophotometer: BioTek, Winoski, USA).

TAC was assessed using the ABTS Antioxidant Capacity Assay Kit (BQC Redox Technology, Spain), following the manufacturer’s protocol. This method is based on the ability of antioxidants present in the sample to reduce the ABTS^●+^ radical cation (2,2’-azino-bis(3-ethylbenzothiazoline-6-sulfonic acid)), leading to a decrease in absorbance at 734 nm. The antioxidant capacity was quantified by comparison with a Trolox standard curve and expressed as Trolox Equivalent Antioxidant Capacity (TEAC), in micromolar (µM Trolox).

Alkaline phosphatase activity was determined using a kinetic colorimetric method with p-nitrophenyl phosphate (pNPP) as substrate. The enzymatic hydrolysis of pNPP to p-nitrophenol was monitored by measuring the increase in absorbance at 405 nm for 10 minutes at 37 °C in a final reaction volume of 250 μL. Enzyme activity was calculated using a molar extinction coefficient of p-nitrophenol (ε = 18,000 L.mol^−1^.cm^−1^) and expressed as units per milliliter (U/mL), where one unit is defined as the amount of enzyme that releases 1 μmol of p-nitrophenol per minute under the assay conditions.

##### b. Spain overwinter 2024

A similar protocol as previously described started on November the 1^st^ 2024, with winter mortality as a response variable. We selected 80 hives and split them into two apiaries, each containing 40 hives. Each colony was randomly assigned to one of the two experimental groups: control (plain syrup) or Na-Mo_2_O_4_-EDTA (syrup supplemented with Na-Mo_2_O_4_-EDTA). Colonies were fed twice (at D0 and D7) with 2L of syrup, either plain or supplemented with Na-Mo_2_O_4_-EDTA at a concentration of 2 mg/L, depending on the assigned group. In early March (day 120), we censused the winter mortality.

### 2.4 Statistical analysis

All data were analyzed in R version 4.2.3 (R Core Team, 2022) using RStudio (version 2024.04.0).

We first used Linear Mixed Models (LMM) to test the treatment effect (control *vs*Na-Mo_2_O_4_-EDTA) on the surface of honey on hive comb and the number of honey bees during the experiments in Greece (2022) and Spain (2023). We used *treatment* and *date* as explanatory variables. The *hive*, or the *hive nested within the apiary* to account for hierarchical experimental design, were used as random factors for the Greek and the Spanish tests, respectively. The response *variable number of honey bees* was log transformed to optimize model fit. We then tested whether the winter mortality differed between the two treatments using a generalized linear mixed model (GLMM) with Binomial error distribution and the logit-link function. We considered the *experiment* (Spain 2023, 2024 and France 2023) and the *colonies nested within the apiary* as random factors. We used the odd ratio (and its 95% confidence interval) to obtain the relative risk (Grant, 2014) as a measure of effect size. An independent two-tailed t-test was applied to test if the honey bee mortality was different between both treatments after the normality of distribution was checked using Shapiro-Wilk test. Finally, we used LMMs to test the treatment effect on both ALP and TAC biomarkers. We set *treatment* and *date* as explanatory variables and the *hive nested within the apiary* as a random factor.

The LMMs and the GLMM were conducted using the lmer or glmer functions respectively from the package ‘lme4’ (Bates et al., 2015). Then, if the model test was significant, Tukey’s Honest Significant Difference Test was used as post hoc analysis with the glht function of the package ‘multcomp’. LMMs and GLMMs were used to account as the dataset involved repeated measures and/or nested variables. Model selection was done according to the Akaike Information Criterion (AIC). Model fits were visually checked using QQ plots and tested for overdispersion with the testDispersion function of the DHARMa package (Hartig, 2019). A significance level of α = 0.05 was used for all tests.

## 3. Results and discussion

The complex Na-Mo_2_O_4_-EDTA, depicted in Figure 1A, has been reported several times in the literature since 1956 (Pecsok and Sawyer, 1956), notably as a biomimetic model of enzymes (Ott et al., 1977). It is built from a molybdic [Mo_2_O_4_]^2+^ core surrounded by an EDTA organic molecule widely used in the food industry, which stabilizes the complex and probably contributes to the bioassimilation of the complex (Fuior et al., 2025).

Several aspects of a product can be evaluated before its commercial distribution in the beekeeping sector, such as the stability of the molecule, its toxicity, and its safety for the honey bee and the hive products. Once these points have been demonstrated, it can be used in beehives under real beekeeping conditions.

### 3.1 Stability of Na-Mo_2_O_4_-EDTA in syrup

ICP-MS studies on aqueous solutions of 1.6 g/L of complex, and on syrup solution at 2 mg/L show that the level of Mo remains constant within 2 years for both of them (Table A1, Appendices). The electronic spectrum measured by UV-Visible spectroscopy in water and in sugar solution constitutes a fingerprint of the complex (Ott et al. 1977, Fuior et al. 2025). Fuior et al. (2025) demonstrated with this technique that the complex is stable even at low concentrations (0.2 µM), below the one used in the beehives (2mg/L, 3 µM). The current study was performed to establish stability throughout the time, for water and sugar syrup solutions, two typical formats which could be proposed for a further commercialization. In case of degradation of the complex, it would decompose into polyoxomolybdate species which would precipitate, and the Mo-content would decrease with time. It is not the case in the study presented here. The complex Na-Mo_2_O_4_-EDTA is stable over time under regular storage conditions (e.g. in a cupboard at room temperature). This point is important for the beekeeping industry.

### 3.2 Safety of honey

To be commercialized, the honey produced by hives fed with Na-Mo_2_O_4_-EDTA must be unaltered and free from residues of the complex Na-Mo_2_O_4_-EDTA, including Mo, EDTA and its decomposition products. This point was assessed by administering to the hives four times the dose we used in the field tests, i.e. 32 mg per hive instead of 8 mg. Even with this high dosage, Mo was not detected in any of the honey samples (Table A2, Appendices). The results of the analysis of EDTA and its main degradation products did not show any traces of the molecules neither in honey from the brood nor from the supers (Table A3, Appendices). Furthermore, honey samples collected in the brood area and in the supers matched the EU regulations in terms of pH, humidity, sugar content, conductivity, HMF content, and Schade degree of Diastase (Table A4, Appendices). The safety of honey from hives fed with Na-Mo_2_O_4_-EDTA was proven, even in case of an overdose of the complex, allowing beekeepers to use it. The main results are gathered in Table 1.

**Table 1.**
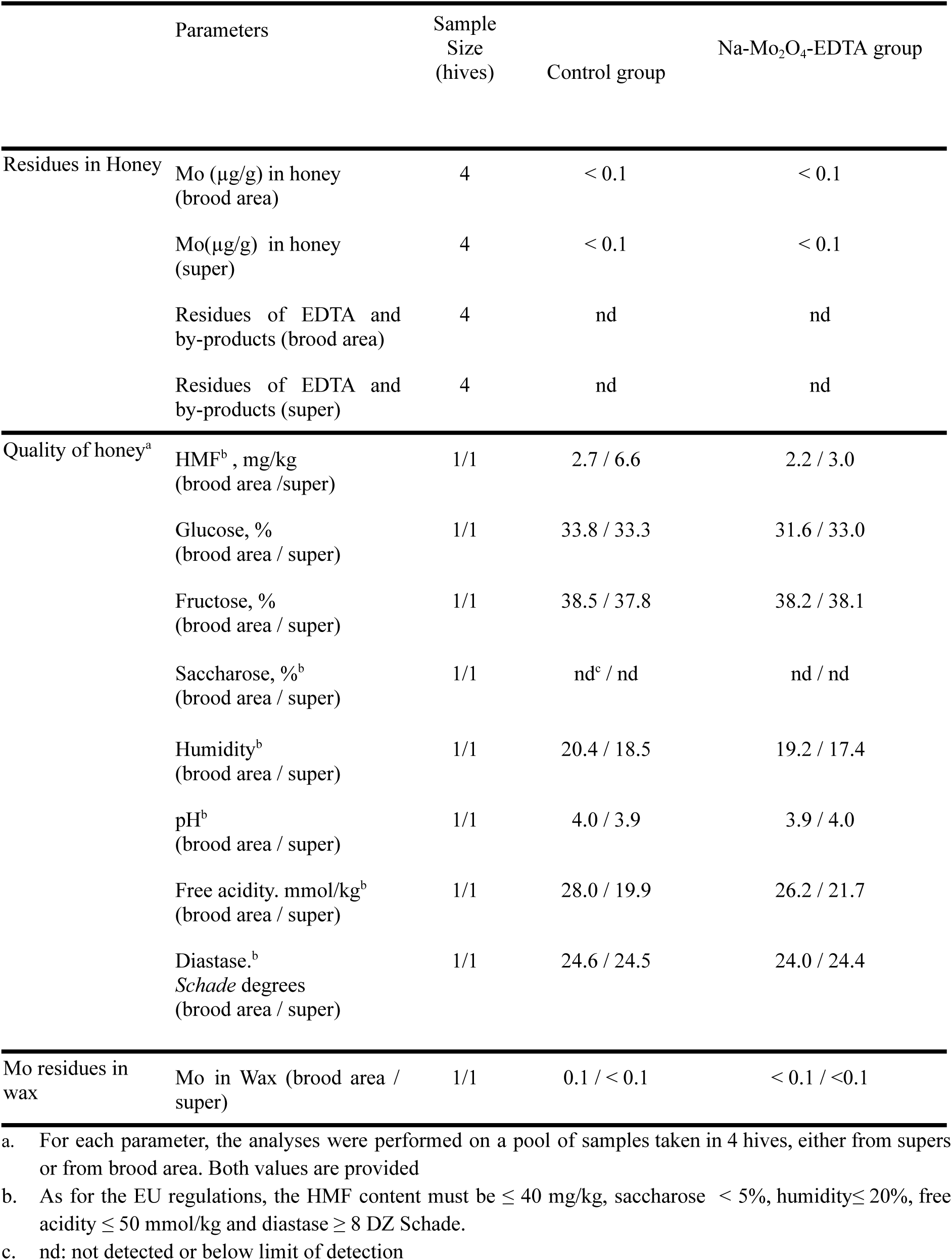
Summary of results of the residue analyses and the quality of honey obtained by feeding two groups of 4 hives with sugar (control) or sugar supplemented with Na-Mo_2_O_4_-EDTA at 32 mg/hive. Raw data and details are given in Appendices (Tables A2, A3, A4).

Besides, in the wax samples, Mo was detected at 0.1 mg/kg in only one pooled sample from the control group and was not detected in any samples from the Na-Mo_2_O_4_-EDTA group (Table 1). This is in accordance with the lipophobic character of the complex (solubility lower than 2.5 mg/L in octanol) *vs* its high hydrophilic character (solubility higher than 1000 g/L in water). We conclude that Na-Mo_2_O_4_-EDTA did not leave any residues in the wax.

### 3.3 Safety of Na-Mo_2_O_4_-EDTA for honey bees

A tolerance study was carried out through the Greek field campaign. The cumulative mortality was 128 bees (±60) and 114 bees (±40) in the control and in the Na-Mo_2_O_4_-EDTA groups respectively (mean ± sd), with no significant difference between both treatments (t-test, p = 0.47). The non-toxicity of the complex Na-Mo_2_O_4_-EDTA has already been established on different organisms and models such as protozoa, mammalian cells, mice, *Daphnia magna*, and honey bees where a Na-Mo_2_O_4_-EDTA dosage of ranging from 8 to 80 mg/hive showed no increase in bee mortality (acute and chronic toxicity, in the laboratory and the field) (Fuior et al., 2025, 2022). The complex Na-Mo_2_O_4_-EDTA is not toxic for many organisms and especially for honey bees either in laboratory conditions or in field conditions.

### 3.4 Impact on honey reserves and honey bee population

Results from the honey reserves, defined as the surface of combs covered by honey in hives from control and Na-Mo_2_O_4_-EDTA groups, are depicted in Figure 2 (A and B). In Spain, we found that the date of assessment had an impact on honey reserves (LMM, χ² = 128.1, p < 0.0001), and we observed no effect of the treatment (LMM, χ² =0.07, p = 0.077) (Figure 2A). At the beginning of the experiment (D0), before winter, honey reserves were similar between hives of the same apiary. In this experiment, honey reserves were not influenced by the feeding with Na-Mo_2_O_4_-EDTA. Conversely, in the Greek experiment, the date of assessment and the treatment had a significant effect on honey reserves (LMM, χ² = 7.01, p = 0.008 and χ² = 7.67, p = 0.005 respectively) (Figure 2B) with an increase of 107% at D56 compared to the control (Tukey HSD, p = 0.025). Based on the results in Greece, it appears that the Na-Mo_2_O_4_-EDTA complex can improve the honey reserves in the supplemented colonies, possibly by triggering or stimulating foraging. Fuior et al. (2025) observed a similar benefit of the complex from a field test in Moldova, with an increase of 49% in honey production. Na-Mo_2_O_4_-EDTA complex might help in the xenobiotics detoxification (see also par. 3.6 below), therefore increasing the health, the longevity and the foraging ability of the bees.

**Figure 2.**
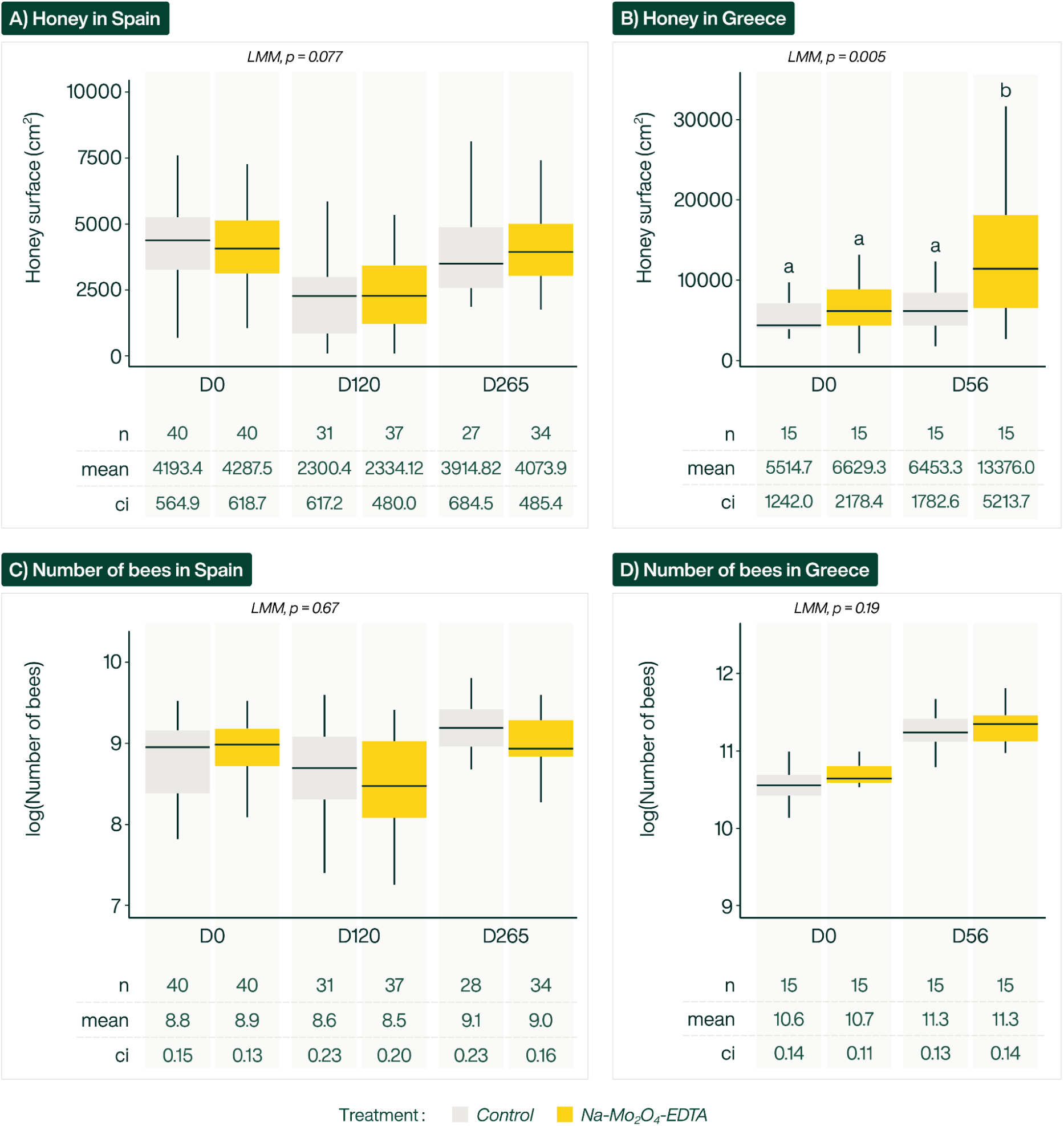
Surface of comb occupied by honey (cm^2^) in hives from control or Na-Mo_2_O_4_-EDTA groups, A) in the Spanish experiment at days 0, 120 and 265 and B) from the experiment in Greece at day 0 and 56. The letters within the bar plot indicate the results of pairwise comparisons between treatments. Number of honey bees (log transformed) in hives from control or Na-Mo_2_O_4_-EDTA groups, C) in the Spanish experiment at days 0, 120 and 265 and D) from the experiment in Greece at day 0 and 56. Different colors represent different treatments. Boxplots show median (horizontal crossbar) and interquartile ranges, the point inside each box indicates the mean. Each table shows the descriptive statistics associated with the boxplot (n: sample size, mean, ci: 95 percent confidence interval of the mean).

Several hypotheses may explain the variability of results between experiments. First, the timeline of feeding was different between both experiments. Then, the inherent differences in beekeeping practices among countries make the direct comparisons between individual apiaries complex. Specifically, in Spain - and particularly in Extremadura - beekeeping often involves a large number of colonies per apiary. Such a scale requires less individualized hive management. In contrast, the Greek apiary was affiliated to a research institute with a smaller number of colonies, allowing for more detailed and individualized management practices. The fact that we were able to detect an effect in the Greece experiment while the sample size was smaller, could have been explained by the use of nuclei with a similar bee population and honey reserve at the beginning of the experiment. This procedure allowed a better control of the intrinsic variability among honey bee colonies. However, the intrinsic variability at day 0 was similar in Spain and in Greece (honey reserves, mean *CV* = 42% and 40% for the control group in Spain and in Greece respectively). The other hypothesis is a variability of response between sites. The weather conditions and available floral resources may explain the variability between apiaries. The period of feeding probably plays a role. The genetic origin of the bee populations was different between the 3 countries, with *A. mellifera iberiensis* in Spain and *A. m. macedonica* in Greece. Bee populations of genetic origin are expected to respond differently to stimulus and manipulations. To overcome this high variability between sites, spreading the investigation in multiple regions with a harmonized protocol will be particularly useful.

Measurements of the estimated number of bees are presented in Figures 2C and 2D. The pattern of the bee population is consistent with what is expected from a colony, with a slight population decrease in winter and then an increase in the number of bees in spring. We found no differences between the two treatments, neither in Spain (LMM, χ² = 0.17, p=0.67) nor in Greece (LMM, χ² = 1.70, p = 0.19). We could have interpreted an increased bee population as a signal of an increase in the egg laying rate of the queen or in the survival rate of each bee caste. From the present study we conclude that Na-Mo_2_O_4_-EDTA has no effect on these two variables.

### 3.5 Winter mortality

The winter mortality was tested within six apiaries, with in total 123 and 124 hives in control and Na-Mo_2_O_4_-EDTA groups respectively. As shown in Figure 3, the winter mortality of the control group depends on the apiary. The mortality rate was around 10% in France in 2023 in both experimental apiaries, while in Spain in 2023 it was about 20-25%, and it increased to 35-42% in winter 2024. This result is coherent with the mortality rate observed in Spain in winter 2019/2020 reaching 36.5% (Gray et al., 2023). The treatment had a significant effect on winter mortality (GLMM, χ² = 4.72, p = 0.029), which significantly decreased for the Na-Mo_2_O_4_-EDTA group compared to the control group (Tukey HSD, p = 0.029). The risk of colony loss was twice lower in the Na-Mo_2_O_4_-EDTA group compared to the control group (odd ratio: 0.455, 95% CI: 0.21-0.91, risk ratio: 0.51). The average mortality decreased by 44,3% in the Na-Mo_2_O_4_-EDTA group. A similar trend was observed in each apiary, with a decreased mortality ranging from 27% to 75% in the Na-Mo_2_O_4_-EDTA groups compared to the control groups (Figure 3). This result is coherent with the results obtained with the lithium salt Li-Mo_2_O_4_-EDTA (Fuior, 2025) in California USA, where a 44% to 80% decrease of the mortality rate was observed.

**Figure 3.**
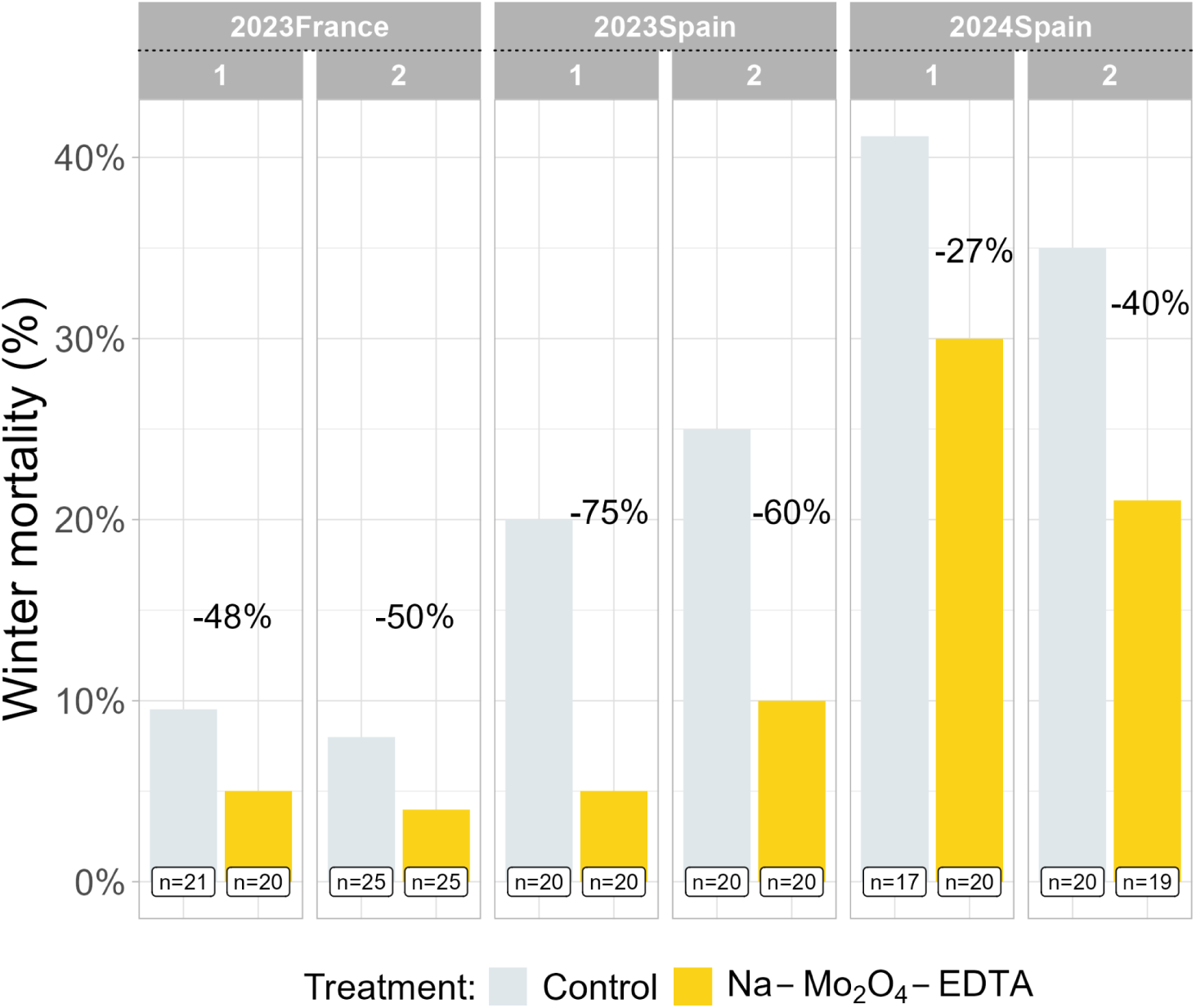
Winter mortality rate of honey bee colonies from each experimental group (control or Na-Mo_2_O_4_-EDTA) for the field campaigns in France (2023) and Spain (2023 and 2024). The % indicate the reduction of the mortality rate in the Na-Mo_2_O_4_-EDTA groups compared to the control groups and n refers to the sample size (number of colonies).

Winter is a critical season for the survival of the honey bees and most colony losses occur during winter. The percentage of losses varies across regions, apiaries, and years, and depends on numerous parameters such as the climate, available resources, and diseases. Population size and honey reserves have been described as colony traits that support honey bee overwintering (Minaud et al., 2024). From our experiments in Spain (feeding in November), we observed that mortality could be further decreased through supplementation with Na-Mo_2_O_4_-EDTA for colonies with a similar number of honey bees and honey reserves (Figures 2A and 2C).

This result is highly relevant to the beekeeping sector affected by high winter mortality. For example, the region of Extremadura frequently experiences a high winter mortality rate (García-Vicente et al., 2024a), while it is an important beekeeping region in Spain (MAPA, 2023). The cost of hive replacement and revenue loss due to high mortality rates put beekeepers under serious economic strain (Bixby et al., 2023). A significant reduction of colonies’ mortality is an important result for the beekeeping industry. Beekeepers often use supplementation to compensate for honey harvest (Papežíková et al., 2020; Quinlan et al., 2023). Protein supplements have been demonstrated to reduce winter mortality in certain conditions (García-Vicente et al., 2024b), while caged tests have evaluated the potential benefits of the trace element zinc (Zhang et al., 2015) or a mixture of chelated transition metals (Ghasemi et al. 2025). However, to our knowledge, this is the first study demonstrating a significant reduction of winter mortality using a trace element based feed supplement through large scale field trials.

Our results suggest that molybdenum might have a role in improving the health of winter honey bees. The hypopharyngeal glands (HPGs) play an important role in honey bee health (Dohanik et al., 2024; Hrassnigg and Crailsheim, 1998). They are notably involved in the production of vitellogenin, which plays an important role to protect bees from oxidative stress and contribute therefore to their longevity (Omholt and Amdam, 2004; Seehuus et al., 2006). Fuior et al. (2025) observed by X-Ray fluorescence that HPGs naturally contain traces of molybdenum. Moreover, in laboratory conditions, the complex Na-Mo_2_O_4_-EDTA is assimilated by honey bees, especially in the brain and in the HPGs. While the role of Mo in HPGs remains unknown, its presence in these glands constitutes one hypothesis to explain the observed effect on the winter mortality of colonies. A better understanding of the causes of winter mortality in the studied region would also improve the understanding of the mode of action of the feed supplement.

### 3.6 Effect of Na-Mo_2_O_4_-EDTA at the biochemical level

To further understand the mode of action of the complex in honey bees, the alkaline phosphatase activity and the oxidative stress in worker bees were studied within the Spanish field test (2023).

Alkaline phosphatase (ALP) is an enzyme located in the midgut of bee that catalyzes the hydrolysis of phosphate substrates, even if its overall role remains unclear. ALP uses zinc and magnesium as cofactors. In other organisms, molybdate and EDTA have been described as competitive and irreversible inhibitors of the ALP (Le-Vinh et al., 2022). For this reason and as the complex Na-Mo_2_O_4_-EDTA is administered through oral feeding, we looked at the potential consequences of the supplementation on the ALP activity during the experiment in Spain. We found a significant effect of both the day of sample collection and the treatment on the response variable (LMM, χ² = 29.9, p < 0.001 for day; χ² = 4.09, p = 0.042 for treatment). In particular, ALP activity increased in the Na-Mo_2_O_4_-EDTA group at Day 120 (Tukey HSD, p = 0.006), before returning to levels comparable to the control group by Day 265 (Tukey HSD, p = 1) (Figure 4A).

**Figure 4.**
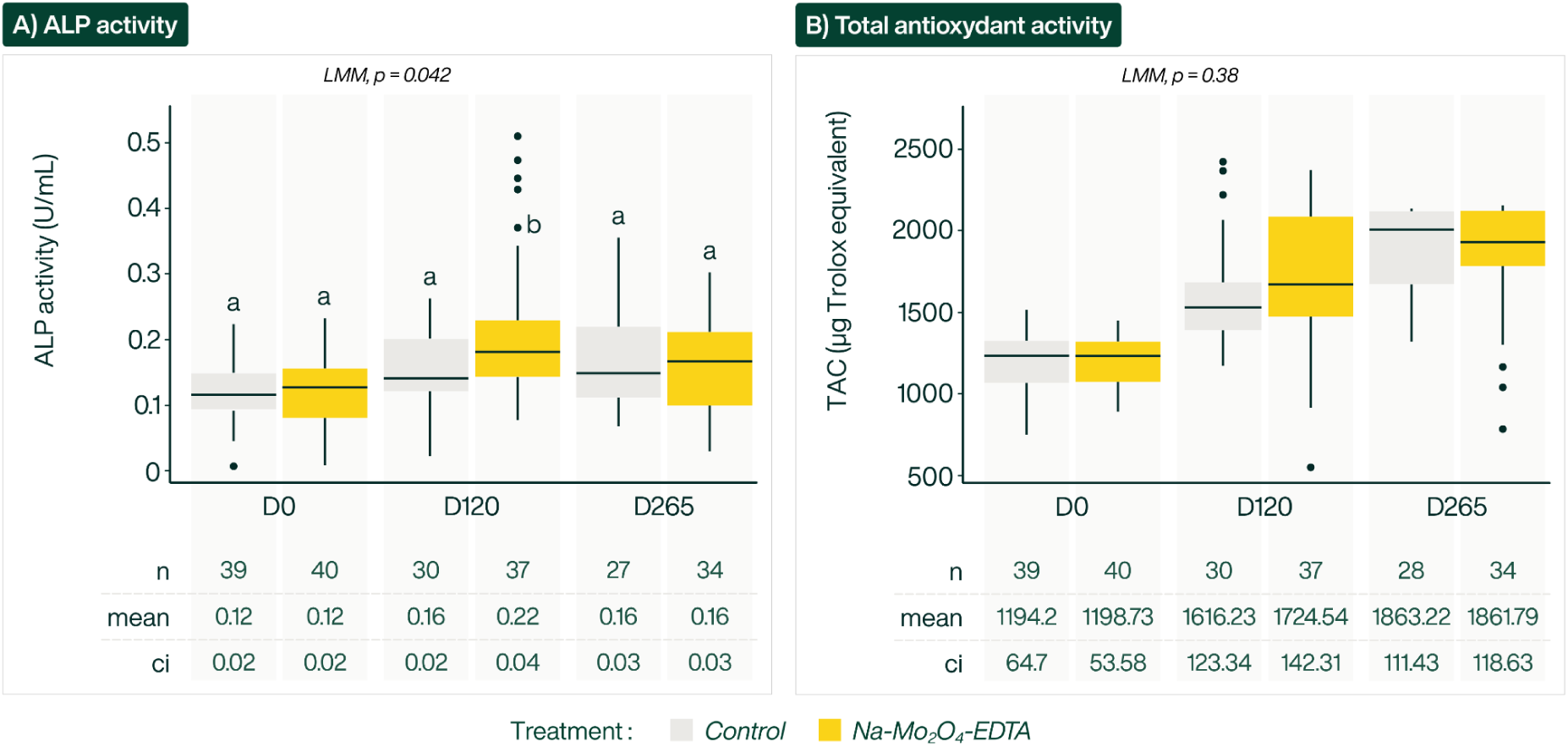
A) Alkaline phosphatase (ALP) activity (U/mL) and B) total antioxidant capacity (TAC) expressed as Trolox Equivalent Antioxidant Capacity (TEAC, µM Trolox) from control and Na-Mo₂O₄-EDTA groups from samples of worker bees from the Spanish experiment at days 0, 120 and 265. Each table shows the descriptive statistics associated with the boxplot (n: sample size, mean, ci: 95 percent confidence interval of the mean).

The consequences of an ALP level modification in honey bees are not clear in the literature. An increase of ALP level of about 20% above the control level has been observed in honey bees exposed to low doses of thiamethoxam under laboratory conditions, and the enzyme has been proposed as a biomarker of exposure to this insecticide (Badiou-Bénéteau et al., 2012). On the other hand, imidacloprid exposure decreased the ALP activity (Paleolog et al., 2020). An activation of the ALP activity has also been observed in bees exposed to 1 mM of copper(II) salts without causing any significant adverse effect (Bounias et al., 1996), while some studies have associated an increase in the ALP to a greater bees’ defense systems and longevity (Strachecka et al., 2014). From our results on Na-Mo_2_O_4_-EDTA non-toxicity for honey bees, we conclude that the observed change in the ALP activity does not jeopardize, and may even benefit honey bee health. Molybdenum acts as a cofactor for several enzymes, for example sulfite oxidase and xanthine oxidase (Hille, 2002). Although the direct involvement of these specific Mo-dependent enzymes in honey bee metabolism have not been fully elucidated, it is plausible that Mo supplementation could indirectly influence overall metabolic flux. In mammals, ALP has been described to enhance immunity functions (Chen et al., 2011). Enhanced availability of Mo might improve the efficiency of certain metabolic pathways, potentially leading to compensatory or adaptive upregulation of ALP to process nutrients more effectively or make bees more resilient to stress factors such as insecticides (Wang et al., 2011). However, the biological relevance of this specific change, and its direct impact on colony health or its productivity, requires a cautious interpretation. Further research is needed to link the temporary increase of ALP activity to measurable improvements in honey bee vitality or specific disease resistance.

TAC is defined as a measure of the combined ability of all antioxidants present to neutralize reactive oxygen species (ROS). While we initially hypothesized that molybdenum supplementation could enhance antioxidant status, notably due to the reduced oxidation state of Mo atoms in the complex, our results did not show significant differences between the treated and control colonies regarding TAC levels. There was a significant effect of the day of sample collection (LMM, χ² = 186.0, p < 0.001) and no significant differences in TAC values between both treatments (LMM, χ² = 0.73, p = 0.38) (Figure 4B). Fuior et al. (2025), demonstrated some potential antioxidant activity in honey bees supplemented under field conditions. We could not confirm this trend in the present study. Several differences in the experimental design can explain why the antioxidant effect could not be reproduced in the present study. The different experimental timeline may explain the difference between the result in Fuior et al. (2025) and the result of the present study. Bees were collected 14 days after supplementation while we collected them at days 120 and 265. The experiments took place in two geographical regions (Moldova and Spain), which may add variability in the observations. Finally, the biological matrix was also different, with the entire bees used in present study and the hemolymph used in the former experiment. Moreover, TAC values in honey bees are known to be highly variable and influenced by multiple environmental and physiological stressors (Nicewicz et al., 2025). This variability likely lowered the statistical power of our experiment and masked any subtle or moderate effect of the supplement.

## Conclusion

The aim of this study was to evaluate the potential benefits of feed supplementation with a molybdenum-based complex for honey bees, as well as its non-toxicity and suitability for the use by commercial beekeepers.

It was first demonstrated that a few milligrams of Na-Mo_2_O_4_-EDTA complex, improved the honey reserves by up to +107% in an apiary in Greece. It was also demonstrated that the quality of the honey produced was not altered even in case of an important dosage of the supplement (32 mg/hive) ensuring consumer’s safety..

The main result of this study is a significant reduction in winter mortality rates, one of the critical issues in the beekeeping sector. Na-Mo_2_O_4_-EDTA diluted in sugar syrup reduced winter mortality by an average of 44% compared to the control group. The study across different climatic and geographical conditions confirmed the consistent mortality reduction, independently of local beekeeping practices and the genetic origin of the bee used.

The mode of action of this molybdenum-based complex is not yet elucidated, which opens a number of hypotheses to investigate.

Although earlier studies demonstrated the complex’s potential as an antioxidant, the current study did not demonstrate any impact on TAC, highlighting the need for additional investigations to clarify its precise mechanisms of action and physiological impact.

Further investigations are also needed to evaluate the effects of the complex on the morphology and the size of the hypopharyngeal glands, on the vitellogenin expression levels, on the quality of the royal jelly or on expression of Mo-based detoxification enzymes such as xanthine oxidase.

Understanding the mode of action of the feed supplement would also benefit from research on the various castes - workers, drones or queens - and the various life stages - larvae or adult - of honey bees.

Finally, it would be interesting to understand if the benefits observed with this complex are due to an environmental deficiency in the molybdenum trace element. Such a project could help to identify the lack of essential metals in honey bee’s organisms and participate in developing new areas for feed supplementation with micronutrient-based complexes. A recent study on the use of a mixture of minerals is in favour with this hypothesis (Ghasemi et al., 2025).

Beyond these considerations, the main takeaway from this study is that the Mo-based complex Na-Mo_2_O_4_-EDTA meets all essential criteria: handling, stability, ease of use and efficacy, positioning it as a valuable tool for the beekeeping industry, especially in regions facing high colony mortality rates.

## Acknowledgments

We sincerely appreciate the assistance of Dr. Diana Cebotari for her help for the experimental part and M. Antoine Stamos for figures editing. The authors wish to thank the professional beekeepers, Pascal and Marie Valois (SAS Val’Api), Frank Alétru (Apidev) and José Luis Díaz Serrano that welcomed the tests in their apiaries. We thank the three anonymous reviewers who provided helpful and constructive comments on the manuscript. Mrs Aneta Ozieranska is gratefully acknowledged for English correction.

## Author contributions

Conceptualization: SF & MH; Data curation: MBC, PC, CR; Formal analysis: CR; Funding acquisition: SF ; Investigation: MBC, EJCV, AP, MM, NH, DR, BP, PC, FH, LC, SMM; Methodology: SF & MH ; Project administration: SF ; Supervision: SF ; Validation: MBC, DR, BP, FH; Visualization: CR & MH; Writing – original draft: CR, MH, SF, SMM; Writing – review and editing: SF & CR

## Funding

The studies in France and Spain 2024 were funded by Oligofeed SAS that aims to commercialize the product based on the research presented in this paper. We would like to thank EIT Food, co-funded by the European Union, for supporting and funding the campaign in Spain 2023 and SATT Paris-Saclay (Project APIMONA) for funding the campaign in Greece.

## Competing interests

Two authors work for Oligofeed SAS (CR & MH) and one is a consultant at Oligofeed SAS (SF). One author is linked to a patent about the technology used in this study (SF, European Patent EP4185594B1 delivered on 4th december 2024). The authors declare these interests in the interest of full transparency and affirm that the reported findings are presented objectively and without bias.

## Data availability

The data from this study has not been deposited into any public repository and will be made available upon request.

