## Appendices for "Reducing Honey Bee Winter Mortality with Molybdenum Supplementation: Field Evidence Across Europe"

**Table A1**. Analysis by ICP-MS of the Mo-content in an aqueous mother solution of complex Na-$Mo_2O_4$-EDTA (1.6 g/L in complex, which translates by an expected value of 465 mg/L of Mo) and a syrup solution of Na-$Mo_2O_4$-EDTA at 2 mg/L, which translates by Mo content expected to be 0.58 mg/L. The concentration of Mo does not vary significantly with time over 2 years.

| **Date** | **Mo content (mg/L)** | |
|---|---|---|
| | Na-$Mo_2O_4$-EDTA 1.6 g/L in water | Na-$Mo_2O_4$-EDTA 2 mg/L in syrup |
| 26/07/2022 | 510 | 0.64 |
| 29/09/2022 | 480 | 0.60 |
| 01/12/2022 | 510 | 0.60 |
| 08/02/2023 | 510 | 0.60 |
| 05/04/2023 | 490 | 0.57 |
| 26/05/2023 | 510 | 0.57 |
| 28/07/2023 | 460 | 0.61 |
| 29/09/2023 | 510 | 0.68 |
| 07/12/2023 | 480 | 0.61 |
| 01/02/2024 | 510 | 0.62 |
| 05/04/2024 | 500 | 0.61 |
| 07/06/2024 | 510 | 0.62 |
| 02/08/2024 | 490 | 0.61 |
| **Average value** | 498 | 0.61 |
| **Expected value** | **465 mg/L** | **0.58 mg/L** |

**Table A2.** Results of Mo contents in honey and wax samples taken from beehives of the experiment performed in west France 2020.

| | Hive ID | Mo in honey from brood area (mg/kg) | Mo in honey from supper (mg/kg) |
|---|---|---|---|
| Control group | 4 | <0.1 | <0.1 |
| | 13 | <0.1 | <0.1 |
| | 16 | <0.1 | <0.1 |
| | 20 | <0.1 | <0.1 |
| Na-$Mo_2O_4$-EDTA | 63 | <0.1 | <0.1 |
| | 64 | <0.1 | <0.1 |
| | 68 | <0.1 | <0.1 |
| | 70 | <0.1 | <0.1 |
| | | Mo in wax from brood area (mg/kg) | Mo in wax from supper (mg/kg) |
| Control | | 0.1 | <0.1 |
| Na-$Mo_2O_4$-EDTA | | <0.1 | <0.1 |

***Limit of quantification : 0.1 mg/kg; expanded measurement uncertainty: Molybdenum: +/- 27%***

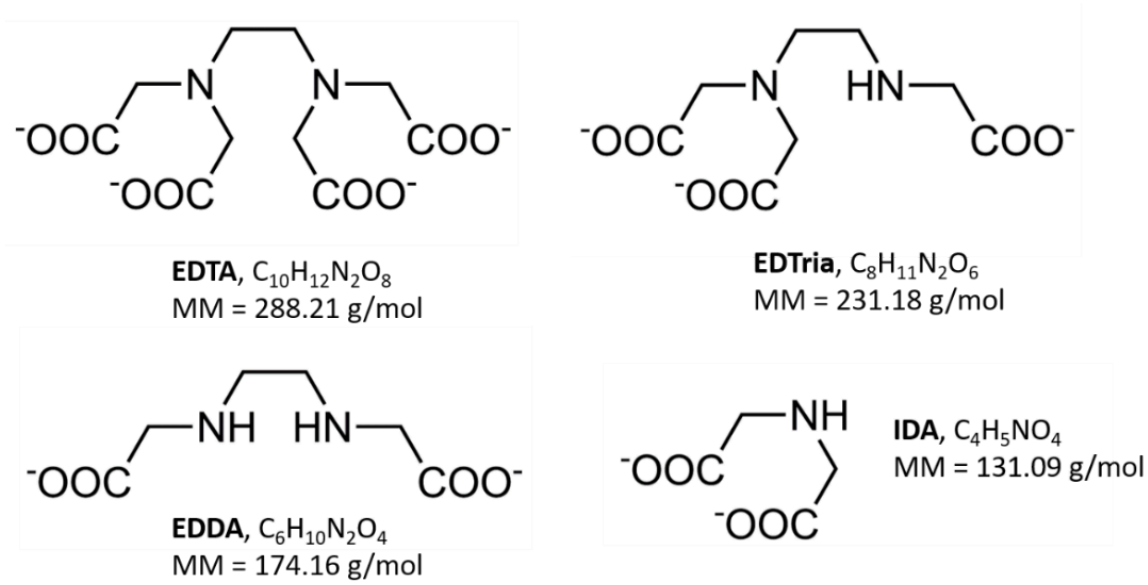


**Figure A1.** Structures of the molecules EDTA. EDDA. EDTria and IDA

**Table A3.** Results of GC-MS analysis of residues of EDTA and its degradation products in honey samples taken from beehives of the experiment performed in west France 2020. n.d. means « not detected at the threshold indicated in brackets (in ppm) »

| Group | Hive | Type | EDTA[a] | EDDA[b] | IDA[c] | EDTria[d] |
|---|---|---|---|---|---|---|
| **Control** | **4** | **Brood** | n.d. (2.12) | n.d. (2.31) | n.d. (2.30) | n.d. (2.24) |
| | | **Super** | n.d. (1.48) | n.d. (1.22) | n.d. (0.92) | n.d. (1.68) |
| | **13** | **Brood** | n.d. (2.59) | n.d. (0.92) | n.d. (1.19) | n.d. (2.45) |
| | | **Super** | n.d. (1.14) | n.d. (1.36) | n.d. (1.11) | n.d. (1.60) |
| | **16** | **Brood** | n.d. (1.73) | n.d. (2.01) | n.d. (0.95) | n.d. (1.27) |
| | | **Super** | n.d. (1.95) | n.d. (0.61) | n.d. (2.28) | n.d. (1.95) |
| | **20** | **Brood** | n.d. (2.04) | n.d. (2.18) | n.d. (1.94) | n.d. (2.19) |
| | | **Super** | n.d. (1.51) | n.d. (2.28) | n.d. (1.34) | n.d. (0.883) |
| **Na-$Mo_2O_4$-EDTA** | **63** | **Brood** | n.d. (2.10) | n.d. (2.43) | n.d. (0.91) | n.d. (0.94) |
| | | **Super** | n.d. (1.39) | n.d. (2.53) | n.d. (1.37) | n.d. (0.62) |
| | **64** | **Brood** | n.d. (1.09) | n.d. (2.33) | n.d. (1.85) | n.d. (0.93) |
| | | **Super** | n.d. (2.29) | n.d. (2.33) | n.d. (0.57) | n.d. (1.84) |
| | **68** | **Brood** | n.d. (2.26) | n.d. (1.90) | n.d. (1.29) | n.d. (2.26) |
| | | **Super** | n.d. (1.73) | n.d. (1.71) | n.d. (2.10) | n.d. (2.15) |
| | **70** | **Brood** | n.d. (0.40) | n.d. (1.84) | n.d. (1.83) | n.d (0.40) |
| | | **Super** | n.d. (1.50) | n.d. (2.21) | n.d. (0.41) | n.d. (0.78) |

a) sum protonated form (CAS No. 60-00-4) and soluble salts. Analyzed as derivative. Extraction of characteristic ions 272 and 415. Retention time: 17.03 min.

*b)* sum protonated form (CAS No. 688-57-3) and soluble salts. Analyzed as derivative. Extraction of characteristic ions 184 and 379. Retention time: 11.81 min.

*c)* sum protonated form (CAS No. 142-73-4) and soluble salts. Analyzed as derivative. Extraction of characteristic ions 211 and 184. Retention time: 9.35 min.

*d)* sum protonated form (CAS No. 5657-17-0) and soluble salts. Analyzed as derivative. Extraction of characteristic ion 397. Retention time: 14.74 min.

**Table A4.** Results of physico-chemical parameters of honey samples taken from beehives of the experiment performed in west France 2020. The honey samples are pooled for each modality by type of honey sample (from brood/body of the hive or from supers).

| **Parameters** | **Control** | | **Na-$Mo_2O_4$-EDTA** | | **Specifications from European union regulations** |
|---|---|---|---|---|---|
| | **Body** | **Honey super** | **Body** | **Honey super** | |
| **Hydroxymethylfurfural. mg/kg** | 2.7 | 6.6 | 2.2 | 3.0 | ≤ 40 mg/kg |
| **Glucose. %** | 33.8 | 33.3 | 31.6 | 33.0 | - |

| | | | | | |
|---|---|---|---|---|---|
| **Fructose. %** | 38.5 | 37.8 | 38.2 | 38.1 | - |
| **Ration F/G** | 1.14 | 1.14 | 1.21 | 1.15 | - |
| **Saccharose. %** | n.d. | n.d. | n.d. | n.d. | <5% |
| **Turanose. %** | 0.9 | 1.2 | 0.8 | 1.1 | - |
| **Maltose. %** | 0.6 | 1.4 | 1.4 | 1.4 | - |
| **Thehalose. %** | n.d. | n.d. | n.d. | n.d. | - |
| **Isomaltose. %** | 0.5 | 0.6 | 0.4 | 0.5 | - |
| **Erlose. %** | 0.1 | n.d. | n.d. | 0.2 | - |
| **Melizitose. %** | n.d. | n.d. | n.d. | n.d. | - |
| **Maltotriose. %** | 0.1 | n.d. | 0.1 | n.d. | - |
| **Humidity. %** | 20.4 | 18.5 | 19.2 | 17.4 | ≤ 20% |
| **Electric Conductivity. mS/cm** | 0.41 | 0.35 | 0.35 | 0.40 | - |
| **pH** | 4.0 | 3.9 | 3.9 | 4.0 | - |
| **Free acidity. mmol/kg** | 28.0 | 19.9 | 26.2 | 21.7 | ≤ 50 mmol/kg |
| **Diastase. *Schade* degrees** | 24.6 | 24.5 | 24.0 | 24.4 | ≥ 8 DZ Schade |

*n.d : not determined because lower than the limit of quantification.*
